# Non-latching positive feedback enables robust bimodality by de-coupling expression noise from the mean

**DOI:** 10.1101/144964

**Authors:** Brandon S. Razooky, Youfang Cao, Alan S. Perelson, Michael L. Simpson, Leor S. Weinberger

## Abstract

Fundamental to biological decision-making is the ability to generate bimodal expression patterns where two alternate expression states simultaneously exist. Here, we use a combination of single-cell analysis and mathematical modeling to examine the sources of bimodality in the transcriptional program controlling HIV’s fate decision between active replication and viral latency. We find that the HIV Tat protein manipulates the intrinsic toggling of HIV’s promoter, the LTR, to generate bimodal ON-OFF expression, and that transcriptional positive feedback from Tat shifts and expands the regime of LTR bimodality. This result holds for both minimal synthetic viral circuits and full-length virus. Strikingly, computational analysis indicates that the Tat circuit’s non-cooperative ‘non-latching’ feedback architecture is optimized to slow the promoter’s toggling and generate bimodality by stochastic extinction of Tat. In contrast to the standard Poisson model, theory and experiment show that non-latching positive feedback substantially dampens the inverse noise-mean relationship to maintain stochastic bimodality despite increasing mean-expression levels. Given the rapid evolution of HIV, the presence of a circuit optimized to robustly generate bimodal expression appears consistent with the hypothesis that HIV’s decision between active replication and latency provides a viral fitness advantage. More broadly, the results suggest that positive-feedback circuits may have evolved not only for signal amplification but also for robustly generating bimodality by decoupling expression fluctuations (noise) from mean expression levels.

## INTRODUCTION

Bimodality is a recurring feature in many biological fate-selection programs (1), such as the HIV active-vs.-latent decision (Fig. 1A). Bimodal expression is a population-wide distribution pattern comprised of two gene-expression modes, each corresponding to a specific fate path (2). The mechanisms that can generate bimodal phenotypes have long been studied and the architecture of underlying gene-regulatory circuits appears to be a key driver of bimodality (3-11). Classically, bimodality has been associated with deterministic bistability in gene circuits (12-15). Deterministic bistability requires ultrasensitive input-output relations and can result from non-linear positive feedback (i.e., Hill coefficient > 1) on a constitutively expressed promoter (16,17). However, many promoters are non-constitutive and instead toggle between inactive and active expression states generating episodic bursts of mRNA production (for review see (18)). The finding that promoters undergo episodic bursts of expression led to a proposal that this toggling alone could generate bimodality without deterministic bistability. Compared to constitutive expression, toggling increases the degrees of freedom in a system [19], and if promoter toggling occurs relatively slowly, the resulting expression bursts can potentially produce bimodality independent of ultrasensitivity (19,20). However, the promoter toggling kinetics required to generate bimodality appeared to be in a small portion of the experimentally observed regime (18,21-23) with experimental measures of intrinsic promoter toggling exhibiting kinetics are typically too fast to produce bimodal expression patterns (Fig. 1B)—specifically, the measured promoter toggling rates where greater than *per capita* protein and mRNA decay rates (18,24,25). Nevertheless, synthetic positive-feedback circuits that slowed toggling could induce bimodality (26). Thus, while computational models showed that promoter ON–OFF toggling was sufficient for bimodal expression (20) and synthetic transcriptional circuits lacking bistable feedback could generate bimodal expression (26), it remained unclear how natural biological circuits exploit this mechanism to generate bimodality without bistability. Here, we examine if promoter toggling can intrinsically generate bimodal distributions in a natural biological system (i.e., human immunodeficiency virus, HIV) and the potential physiological relevance.

**Fig 1.**
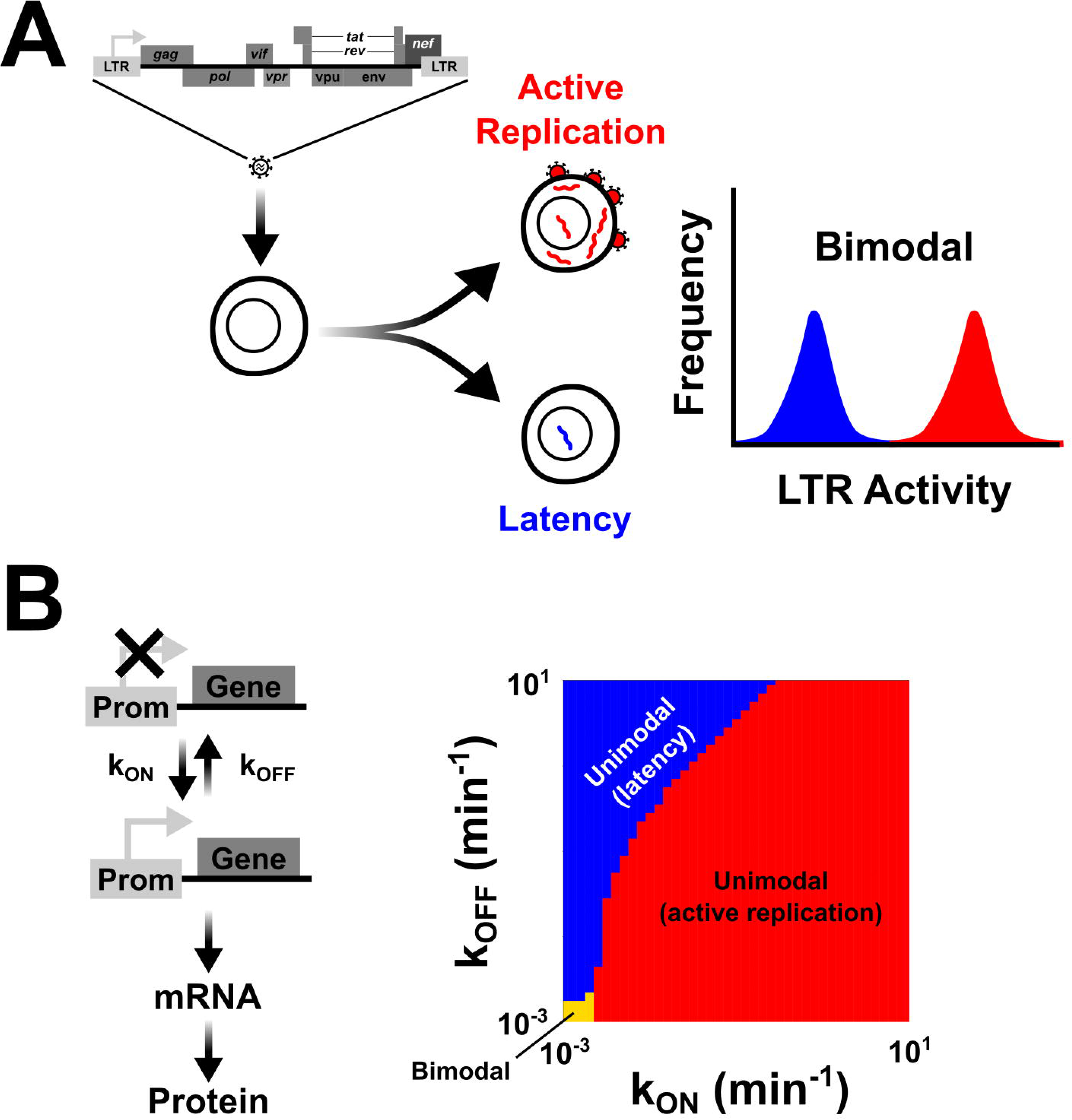
The mechanistic problem underlying bimodal fate-selection programs: promoter toggling is theoretically sufficient to generate bimodality, but only in a narrow parameter regime. (A) A simplified fate-selection decision in HIV. Upon infection of a CD4^+^ T lymphocyte, HIV either enters into an active state of replication (red), producing viral progeny and destroying the host cell or enters into a quiescent state of silenced gene expression termed proviral latency (blue). This fate bifurcation between active replication and latency is not controlled by the cell state (27) but rather by an HIV gene-regulatory program that can generate bimodal gene-expression distributions from its LTR promoter. (B) The LTR is accurately described by a two-state promoter model (e.g., random telegraph models) where the LTR toggles between an inactive (represented by Prom-Gene that is crossed out, top) and active (represented by Prom-Gene) state of expression at rate k_ON_. In the active state, mRNAs are produced, before the promoter flips back to inactive at rate k_OFF._ As a promoter toggles between these active and inactive states, they can produce bimodal distributions in gene-expression products but only within a restricted regime of phase space. Each parameter set was checked to see if it generated unimodal latency (blue), unimodal active replication (red) or bimodality (orange) as described in the materials and methods. For the modality analysis, each mode was required to contain at least 0.1% of the population, otherwise the parameter set was determined to produce a unimodal population.

We focus on HIV as a physiological model system for expression bimodality driving a decision-making process (Fig. 1A). Upon infection of a CD4^+^ T lymphocyte, HIV undergoes a fate-selection decision, either actively replicating to produce viral progeny and destroy the host cell or entering a long-lived quiescent state called proviral latency (28,29). A viral gene-regulatory circuit is both necessary and sufficient to drive HIV fate selection (10). At the core of this decision-making circuit is a virally-encoded transcriptional positive-feedback loop comprised of a single HIV protein, the transactivator of transcription (Tat) that amplifies expression from the virus’s only promoter, the long terminal repeat (LTR) promoter. Molecularly, this positive-feedback loop functions because the LTR is a relatively weak promoter, in the absence of Tat, with RNA polymerase II (RNAPII) elongation stalling ~69-nucleotides after initiation (30). Tat transactivates the LTR by binding to a short ~69-nucleotide RNA-hairpin loop called the Tat-Activation RNA (TAR) loop and recruiting the positive Transcriptional Elongation Factor b (pTEFb)—principally composed of CDK9 and cyclinT1—which hyper-phosphorylates the carboxy-terminal domain (CTD) of RNAPII, thereby relieving the RNAPII elongation block (30,31). Thus, Tat acts much like a bacterial anti-terminator enhancing transcriptional elongation rather than initiation.

Importantly, minimal LTR-Tat positive-feedback circuits are sufficient to generate bimodal expression patterns (32) and in the full-length viral context, this circuit is both necessary and sufficient to drive HIV’s active-vs.-latent decision (27). There are two specific quantitative features of the Tat-LTR feedback circuit that are curious given its obligate role in viral fate selection. First, unlike many other positive-feedback circuits that control phenotypic decisions (33,34), the Tat positive feedback loop is *non-cooperative* (Hill coefficient ≈ 1) and not deterministically bistable (35). Second, the LTR promoter itself displays large episodic expression bursts toggling between ON and OFF states at virtually all integration sites throughout the human genome (24,36,37), raising the possibility that the LTR itself may be sufficient to generate bimodal expression patterns independent of Tat feedback.

In this study, we construct minimal circuits to examine if the LTR itself is capable of generating bimodal expression patterns in the absence of Tat feedback, and then computationally examine the precise role of Tat positive feedback in bimodality. The results indicate that the LTR is intrinsically capable of generating bimodal ON-OFF expression even in the absence of feedback, but that Tat feedback shifts and expands the regime of LTR bimodality into physiological ranges by slowing LTR toggling. In fact, the architecture and parameters of the Tat circuit appear optimized to robustly generate bimodal expression. Given the rapid evolution of HIV, the presence of a circuitry that appears optimized to slow promoter toggling and generate bimodality may be consistent with the hypothesis that the circuit has been selectively maintained and that bimodal expression (between active replication and latency) provides a viral fitness advantage (38).

## RESULTS

### LTR promoter toggling is capable of generating bimodality in the absence of feedback

Previous studies demonstrated that Tat positive feedback can generate bimodal expression patterns from the HIV LTR (32). However, given the large, episodic bursts of expression that characterize LTR activity (24,36,37), we set out to test if the LTR was capable of bimodal expression, even in the absence of feedback (i.e., whether feedback was dispensable for bimodality, possibly having an orthogonal function in HIV). Analysis of experimental and computational literature reports indicated that the regime for generating bimodality through promoter toggling alone fell outside the experimentally observed values of LTR toggling but that slightly slower LTR toggling transitions might generate bimodality without feedback (Figs. 1 and S1).

To test this prediction that Tat feedback was dispensable for bimodality, HIV circuitry was refactored to split the Tat positive-feedback loop (27) into open-loop parts (Fig. 2A). This minimal circuit system allows Tat concentrations to be modulated by doxycycline, and Tat protein stability to be tuned through Shield-1 addition (27). As Tat is fused to Dendra, the Tat concentrations can be quantified, while LTR activity is simultaneously tracked in single-cells. This open-loop doxycyline-inducible circuit was integrated into T cells by viral transduction and cells were exposed to varying concentrations of activator (Dox) and Tat proteolysis inhibitor (Shield-1) –generating ~48 unique unimodal Tat inputs to the LTR (Figs. S2 and S3). Expression profiles from the LTR are all unimodal in the absence of Tat (Figs. S2), in agreement with previous findings (32,36,37). However, in striking contrast, the presence of Tat induces bimodality from the LTR despite the lack of cooperativity or feedback in this open-loop system (Figs. 2B, S2 and S3). In other words, despite a fixed, unimodal concentration of active Tat transactivator, bimodal LTR distributions can be generated, and single-cell time-lapse microscopy confirms that the activity of the LTR is dependent on Tat input (Fig. S4). Based on the known requirements for bimodality to arise from a toggling promoter (Fig. 1), the data suggest that LTR toggling becomes sufficiently slow in the presence of Tat to produce bimodal expression patterns, even in the absence of positive feedback.

**Fig 2.**
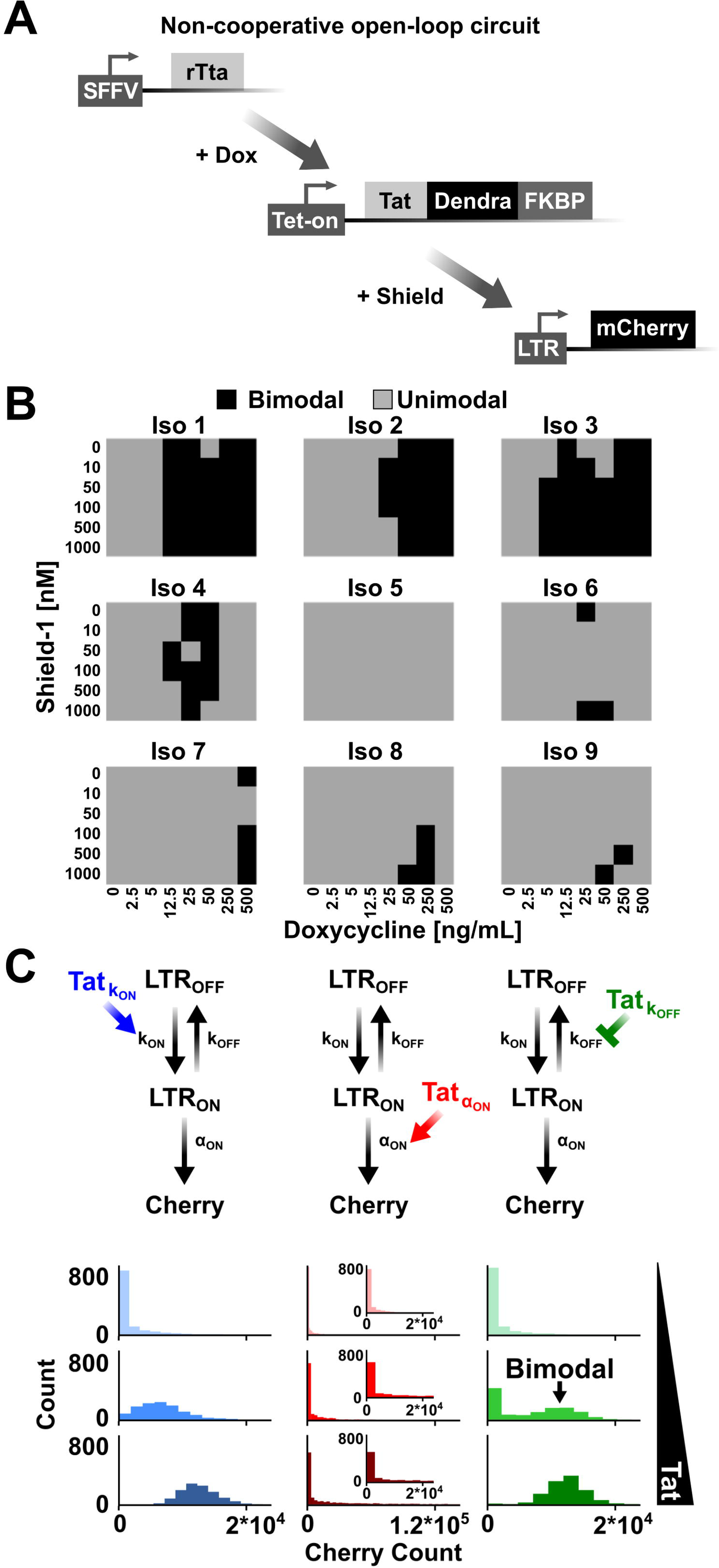
LTR promoter toggling is sufficient to generate bimodality and control HIV fate. (A) Schematic of the open-loop HIV circuit. Doxycycline addition induces transcription from the Tet-ON promoter. Shield-1 addition controls the stability of the Tat-Dendra-FKBP fusion protein. Tat induces transcription from the HIV LTR. (B) The (Iso) term represents an independent isoclonal population so each cell within a clone have the same integration site for the LTR. 9 Iso populations were exposed to 48 different doxycycline and Shield-1 conditions (Figs. S2 and S3) and bimodality was tested for by the Hartigan Dip Test (39) (threshold for determining bimodality was p<0.3, agreeing with an independent test, Fig. S3). Gray squares are determined to be unimodal, black squares are bimodal. (C) Open-loop stochastic model of Tat transactivation of the LTR by one of three mechanisms. Left column, increasing burst frequency by promoting transitions into an active state (increasing k_ON_, blue); middle column, increasing burst size by increasing transcriptional efficiency (increasing α, red); right column, increasing burst size by inhibiting transitions into the inactive state (inhibiting k_OFF_, green). Model equations and details are presented in (Table S1-3). Plotted histograms are steady-state results of 1,000 simulations (at 1000 hours) showing that slowing promoter toggling by inhibiting transitions into the active state is sufficient to generate bimodal distributions (i.e., right column, middle panel). Insets: Zoom of α modulation so the scale of the x-axis matches the k_ON_ (left column) and k_OFF_ (right column) modulation graphs.

### Independent of feedback or cooperativity, LTR promoter toggling is sufficient to control full-length HIV fate

The bimodality in the minimal open-loop system (Fig. 2) represents the two fate paths of the virus—active replication, and proviral latency (40)—and suggested that positive feedback may also be dispensable for controlling viral fate in full-length HIV. Importantly, results from a Tat-deficient full-length HIV virus (27), where Tat is introduced *in trans* (Fig. S5) confirm that Tat feedback is not required to select between alternate HIV fate paths. Thus, unlike other decision-making circuits (17,26), fate selection can occur independent of positive feedback or cooperativity in HIV.

### Tat slows promoter toggling by inhibiting LTR ON-to-OFF transitions, leading to bimodality

To understand the molecular mechanisms enabling LTR bimodality in the absence of feedback, we used a previously validated computational model of HIV (27) and adapted it to an open-loop system where Tat would either modulate: (*i*) burst frequency alone, kON modulation, (*ii*) burst frequency and burst size, k_OFF_ modulation, or (*iii*) burst size alone by affecting transcriptional efficiency, α modulation (Fig. 2C, top, and Tables S1-3). The model results are consistent with previous findings that bimodality is not induced through frequency modulation of the LTR (i.e., k_ON_ modulation) or increases in burst size through transcriptional efficiency, α (24,36,37). However, the model shows that slowing toggling kinetics, or increasing the dwell time in the ON state (i.e., k_OFF_ modulation), is required for bimodality, and if Tat only affects a single parameter, k_OFF_ modulation is necessary and sufficient (Figs. 2C, bottom, and S6).

The interpretation of these results is that while natural LTR promoter toggling is too quick to generate large enough expression fluctuations for bimodality, Tat transactivation is able to slow the kinetics of toggling, expanding the bimodal regime (Fig. 1). The slowing of toggling kinetics reinforces the findings that Tat stabilizes transient pulses of expression from LTR fluctuations (40), thereby effectively reducing k_OFF._ If Tat does stabilize pulses of expression to control gene-expression variability, then the prediction is that altering Tat-feedback strength would, similar to the open-loop system, control the shape of the gene-expression distribution and bimodality.

### Positive-feedback strength controls whether the expression distribution is unimodal or bimodal in HIV

To test the prediction that Tat-feedback strength shapes the expression distribution, we used a synthetic Tat circuit (27) where positive-feedback strength could be manipulated pharmacologically by addition of a small-molecule, Shield-1, that stabilizes Tat proteolysis (Fig. 3A). In this system, a subset of isoclonal cell populations carrying this synthetic circuit naturally generate bimodal distributions (Fig. 3B). These clonal differences are mainly due to the genomic location of HIV integration, which can dictate the transcriptional bursting parameters, and the effectiveness of Tat transactivation (24,36). Though the differences in Tat transactivation potential are not clear, studies have shown that transcriptional parameters of the LTR in the absence of feedback vary due to promoter methylation status, nucleosome acetylation and methylation state, or gene-proximity dependencies (41). When positive-feedback strength is increased, a significant fraction of cells generate bimodal distributions and even convert from a unimodal (low peak) into a bimodal distribution (low and high peak) or from a bimodal (low and high peak) to a unimodal (high peak) distribution (Figs. 3B and S7). In other words, a fraction of isoclones can naturally generate bimodal distributions with low feedback strength (e.g., [0 nM Shield-1]), but with intermediate positive-feedback strength bimodal distributions are more prevalent (e.g., 100 or 500 nM Shield-1).

**Fig 3.**
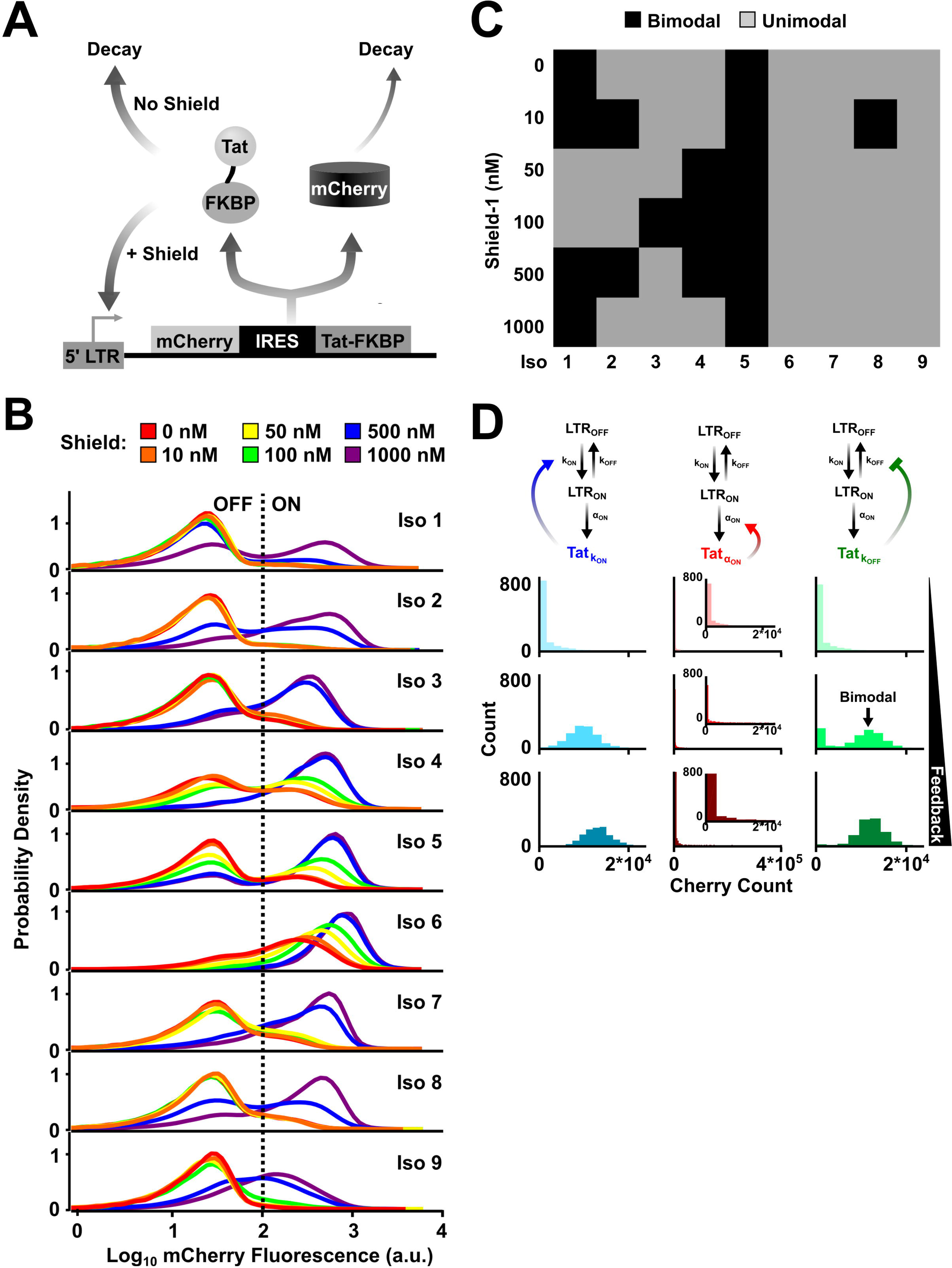
Positive-feedback strength controls whether the expression distribution is unimodal or bimodal in HIV. (A) Schematic of the LTR-mCherry-IRES-Tat-FKBP closed-loop, positive feedback circuit. Tat stability is tuned through the addition of Shield-1 to alter Tat feedback strength, i.e. loop transmission (Fig. S9). (B) Flow cytometry histograms showing bimodal distribution for nine isoclonal cell lines exposed to various concentrations of Shield-1. A fraction of isoclones can naturally generate bimodal distributions with low feedback strength (e.g., red [0 nM Shield-1]), but with intermediate positive-feedback strength bimodal distributions are more prevalent (e.g., green [100 nM Shield-1] or blue [500 nM Shield-1]). The ‘ON’/‘OFF’ threshold was set based on the background level of expression from a naïve Jurkat cell line. (C) Measurement of bimodality for each Shield-1 conidition for each isoclonal population in (B) as quantified by the Hartigan Dip Test. The results agree with another metric for measuring bimodality (Fig. S7). Gray squares are determined to be unimodal, black squares are bimodal. (C) A closed-loop stochastic model (in contrast to the open-loop model in Figure 2C) of LTR promoter toggling that incorporates Tat positive feedback through one of three alternate mechanisms (as in Figure 2C). The steady-state results for 1,000 simulation runs (modeled for 1000 hours) show that Tat inhibition of promoter turn-off is sufficient to generate bimodalities (right column, middle panel), whereas alternate Tat positive-feedback mechanisms are unable to generate bimodality in the requisite parameter regimes.

Importantly, simulations of Tat positive-feedback circuitry corroborate this phenomenon of bimodal expression at intermediate feedback strength if Tat acts by decelerating LTR toggling kinetics (Fig. 3C), in agreement with simulations of the open-loop circuit (Fig. 2). Thus, these simulations indicate that Tat-feedback strength likely alters the natural LTR toggling kinetics set by the local integration site (42) to control HIV bimodal-expression patterns. To test if Tat feedback in fact extends pulses of expression (i.e., *k*_OFF_ reduction) HIV gene-expression was activated to a high-expression state, using tumor necrosis factor alpha (TNFα), and the circuit then allowed to relax back to the unperturbed state under varying feedback strengths. TNFα enhances HIV expression by stimulating recruitment of a p50-RelA heterodimer to NFkB binding sites within the LTR (42). Cells were exposed to TNFα for 24 hrs then allowed to relax back in the presence of strong or weak feedback (Fig. S8). The results show that increasing feedback strength, by dosing cells with increasing amounts of Shield-1, increases the transient in the expressive state, leading to slower transitions from ON to OFF states (Fig. S8), which corroborates previous findings (27,40). Thus, relaxation to various baseline states are dictated by feedback acting on promoter toggling.

One simplifying assumption in the model is that Tat only modulates a single bursting parameter. To test how relaxing this assumption affects bimodal generation, new simulations where Tat could modulate multiple bursting parameters was performed. The models allow Tat to alter both burst size and frequency through either k_ON_ and k_OFF_, k_ON_ and α, or k_OFF_ and α modulation (Fig. S9). Interestingly, the simulations show that any combination of parameters could yield bimodality (Fig. S9). In each scenario, Tat positive feedback yields non-exponentially distributed ‘OFF’ times, and slows toggling kinetics. This result is in agreement with the previous findings that slowing promoter toggling kinetics yields bimodal distributions (Figs. 1-3 and S6).

A few alternate explanations are possible for the observed bimodality. The first is that the bimodality may arise from deterministic cell-to-cell variability (43) where the transcriptional parameters vary between cells, leading to bimodality. However, these minimal circuits display a high level of ergodicity (24,40), suggesting cell-to-cell variability in transcriptional parameters is minimal. Second, HIV feedback may be bistable (i.e. exist in one of two stable states (high or low) (17)). Bimodality observed from bistable circuits results from fluctuations around latching feedback strengths (Fig. S10). Previous studies analyzing fluctuations in noise to measure feedback strength, cooperativity in feedback, or stability of the ‘ON’ state found that HIV feedback lacks the canonical features of bistability (34,35,40). Last of all, HIV feedback may latch, meaning small incrases in Tat would be drastically amplified to saturable levels where the system would then latch in a high state. Note that the latching behavior can be present in deterministically monostable feedback (40). To test this, here, we directly quantified the feedback strength—to test if the feedback-induced bimodality results from latching feedback—by use of the small-signal loop gain, a direct measure of feedback strength (40,44,45). The small-signal loop gain was quantified by measuring changes in LTR expression associated with changing Tat stability (Fig. S10) or increasing Tat concentration (Fig. S11). In agreement with other measures of HIV feedback strength (40), Tat positive feedback appears to be non-latching (Figs. S10 and S11). Interestingly, unlike systems that latch, non-latching feedback strength inherently renders the system relatively insensitive to small fluctuations (46), i.e. HIV will not drastically change expression profile or latch in response to a small fluctuation, lending a molecular explanation for the insensitivity of HIV circuitry to external cues (47,48).

### HIV Tat positive feedback appears optimized to robustly generate bimodal distributions

The combination of non-latching feedback coupled to a toggling promoter allows for bimodal generation across a wide range of Tat concentrations (Fig. 2) and feedback strengths (Fig. 3). Promoters driving non-latching feedback can exhibit extended, transient pulses of expression before reverting back to the initial system state (8). To test if this mechanism of extended-duration transient pulses was responsible for generating bimodality in the LTR, we built a specific model of the LTR to map out the phase space of feedback strengths that would allow for LTR bimodality given the known toggling parameters (Table S1). The model specifically considers promoter toggling coupled to weak positive feedback and examined the effect of changing feedback strength (from weak non-latching to strong non-latching). In agreement with previous theoretical predictions (19,20), intrinsic slow promoter toggling is sufficient to generate bimodality, but only in a very narrow parameter regime (Figs. 1 and 4A).

**Fig 4.**
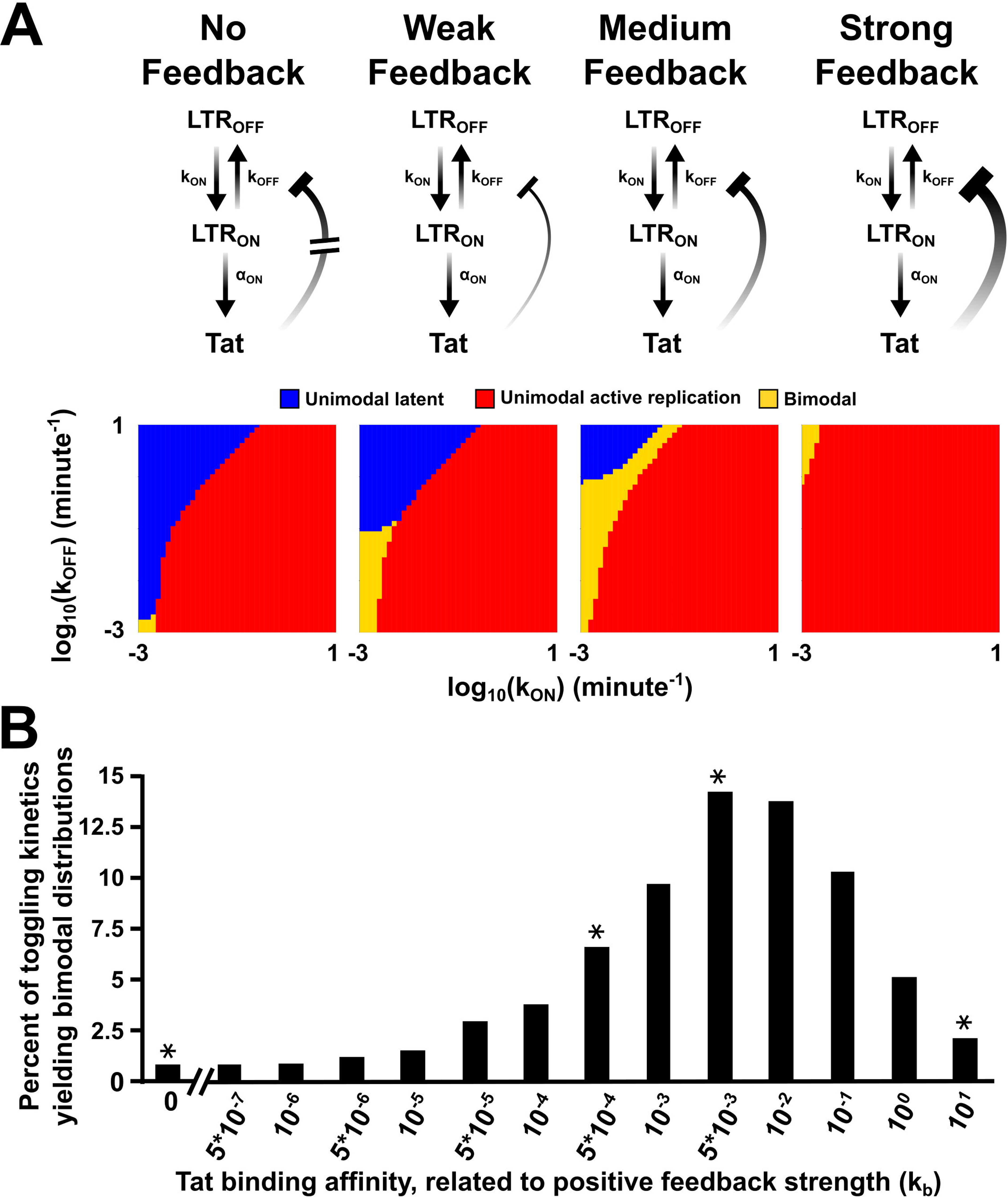
HIV positive feedback appears optimized to robustly generate bimodal distributions. (A) Varying positive-feedback strength changes the toggling kinetics to yield a larger regime for bimodality within the physiological parameter range. The results for the parameter scans are shown for ‘No Feedback’ (left), and increasing feedback strengths. Whether a population was unimodal latent (blue), unimodal active replication (red), or bimodal (orange) was determined for each set of parameters as described in the materials and methods. For the modality analysis, each mode was required to contain at least 0.1% of the population. (B) The percent of toggling kinetics that yield bimodal distributions for varying feedback strengths. The asterisks above the bars represent the feedback strengths shown in (A).

To explore if weak non-latching positive feedback might explain the robust generation of bimodality that was experimentally observed, we incorporated dose-response data from the open-loop circuit into the model and generated an input-output function (Fig. S12) to quantify the relationship between Tat and *k*_OFF_ values. This approach allows of the open-loop data to be mapped unto a model containing feedback (Fig. 4A). The output of the resulting model shows a striking dependence of bimodality on feedback strength (Fig. 4B). Specifically, as feedback strength increases from zero, the bimodality regime significantly expands. However, as feedback increases further, to strong non-latching feedback strengths, there is a drastic reduction in the potential for bimodal generation (Figs. 4B and S13). This acute contraction of the bimodal regime likely results from drastic amplifications of small noise spikes that drive the system to stay on (17). Interestingly, the model predicts that bimodality is generated across ~13% of parameter values for the HIV system (Fig. 4B), in agreement with experimentally observed frequencies for spontaneous bimodal generation across the HIV-integration landscape (32). Thus, HIV’s moderate feedback strength (Figs. S10 and S11) appears optimized to slow promoter-toggling kinetics into the regime enabling bimodality.

### Robust bimodality results from positive feedback decoupling expression noise from mean levels

Since the circuit’s bimodality is ultimately dependent upon fluctuation-driven (i.e., stochastic) extinction of Tat, we next sought to determine how increasing expression levels influenced bimodality. In the classical Poisson or super-Poissonian transcriptional burst models (49), the expression mean scales with variance (σ^2^ ∝ μ) such that the noise magnitude (CV^2^ = σ^2^/μ^2^) decreases proportional to the inverse of the mean squared (Fig. 5A) and the extinction probability can be shown to be (Text S1):

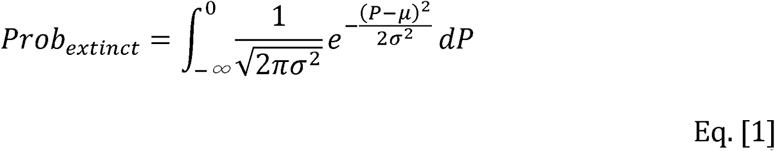

However, non-latching positive feedback breaks the Poissonian relationship such that σ^2^ ∝ μ^N^ with 1 < *N* < 2 (44). In the extreme case where *N* = 2, CV^2^ becomes independent of the mean and the extinction probability becomes (Text S1):

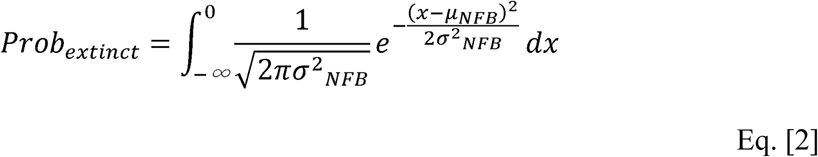

where *σ^2^_NFB_* and *μ_NFB_* are variance and mean for the non-feedback case, respectively. Importantly, Eq. [2] shows that stochastic extinction can be decoupled from the mean (when *N* = 2) and simulations verified that such perfect decoupling was possible (Fig. 5B). Analysis of the experimental data in Fig. 3 shows that the Tat circuit displays partial decoupling of noise and mean with *N* ≈ 1.5 (Fig. 5C). Thus, Tat circuitry enables greater stochastic extinction over a broader range than other circuitries (e.g. no feedback or latching positive feedback) would be able to achieve.

**Fig. 5.**
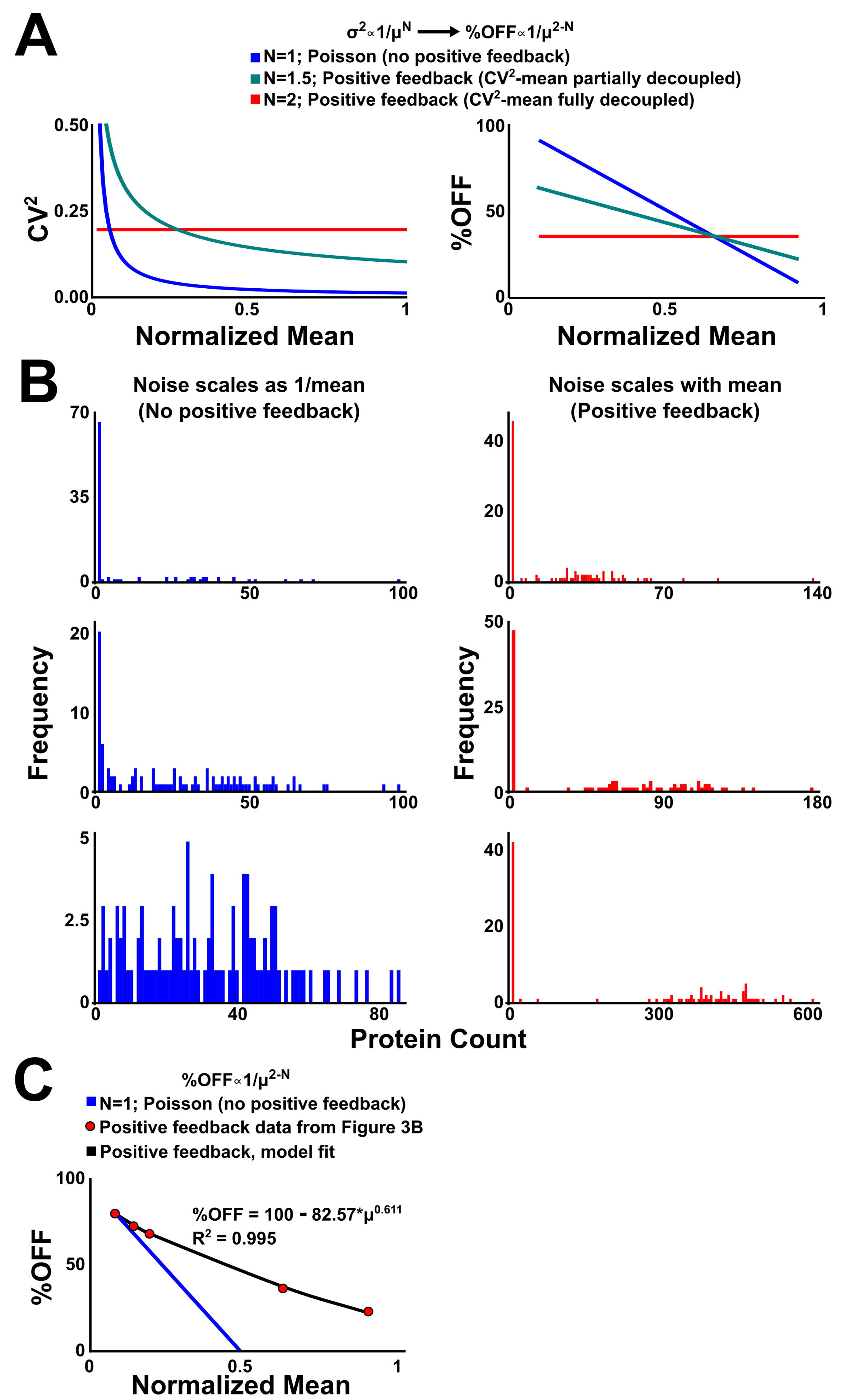
Non-latching positive feedback substantially dampens the Poissonian noise-mean inverse relationship allowing stochastic extinction despite increasing mean-expression levels. (A) In the classical Poisson or super-Poissonian transcriptional burst models (49), the expression mean scales with variance (σ^2^ ∝ μ) such that the noise magnitude (CV^2^ = σ^2^/μ^2^) decreases proportional to the inverse of the mean. Non-latching positive feedback breaks the Poissonian relationship such that σ^2^ ∝ μ^N^ with 1 < *N* < 2 (44). In the extreme case where *N* = 2, CV^2^ becomes independent of the mean. (B) Monte-Carlo (Gillespie) simulations showing that stochastic extinction can be decoupled from the mean (when *N* = 2). (C) Analysis of the data in Fig. 3 shows that the Tat circuit displays partial decoupling of noise and mean (*N* ≈ 1.5).

## DISCUSSION

In summary, HIV’s Tat circuit seems particularly well suited for generating bimodal expression patterns and alternate single-parameter mechanisms for Tat function (e.g. increasing burst frequency alone rather than slowing toggling kinetics) appear to severely limit or completely abrogate the potential for bimodality. The precise architecture of this robust bimodal-generator circuit in such a rapidly adapting virus suggests that bimodality in HIV expression (i.e., latent and active replication modes) may be a beneficial trait that has been selectively maintained (38). In contrast with other known roles for positive feedback (e.g., bistability, noise amplification), these findings demonstrate a further role for positive feedback as a mechanism for robust generation of bimodality (50). On a conceptual level, this ability of positive feedback to expand the bimodal regime into physiological ranges maybe related to positive feedback’s ability to expand the regime where sustained oscillations occur (51,52). Consequently, positive-feedback circuits may have evolved not only for signal amplification but also to stabilize certain dynamic phenotypes (e.g., bimodality and oscillations) in diverse biological systems.

From a basic HIV biology standpoint, these results on Tat’s mechanism of action may have therapeutic implications for HIV cure approaches. Specifically, it has long been known that Tat protein addition reactivates HIV latency more potently than current chromatin remodeling latency-reversing agents (LRAs) such as histone deacetylase inhibitors (HDACi’s) (42). However, there has been no mechanistic explanation as to why Tat protein is more potent than LRAs (e.g., HDACi’s) and the phenomenon was thus considered by some to be an off-target effect. Conventional LRAs (e.g. PKC agonists and HDACi’s) only affect k_ON_ and we have previously shown that agents that simultaneously reduce k_OFF_ and k_ON_ potentiate reactivation (53). Hence, the finding herein that Tat alters k_OFF_, coupled with the magnitude of the Tat-induced k_OFF_ change, provides a mechanistic explanation as to why Tat is so effective for latency reversal. The findings also suggest that Tat-based strategies and conventional LRA strategies could be used synergistically, and new approaches aimed at simultaneously reducing k_OFF_ and increasing k_ON_ would be optimal for ‘shock-and-kill’ strategies, while conversely increasing k_OFF_ and decreasing k_ON_ would be optimal for ‘block-and-lock’ strategies.

## MATERIALS AND METHODS

### Molecular Cloning Procedures

The sequence of Tat from recombinant clone pNL4-3, GenBank: AAA44985.1, M19921, was used. To clone the LTR-mCherry-IRES-Tat-FKBP construct d2GFP was swapped with mCherry using BamHI and EcoRI restriction sites (27). To clone the Tet-Tat-Dendra-FKBP plasmids, Tat-Dendra or Tet-Tat-Dendra was swapped with YFP-Pif from the pHR-TREp-YFP-Pif plasmid (a gift from Wendell Lim’s Laboratory at UCSF) using BamHI and NotI restriction sites. The full-length virus was generated as previously described (27).

### Recombinant virus production and infections

Lentivirus was generated in 293T cells and isolated as described (32,54). To generate the isoclonal closed-loop circuit populations, lentivirus was added to Jurkat T Lymphocytes at a low MOI to ensure a single integrated copy of proviral DNA in infected cells. Cells were stimulated with tumor necrosis factor alpha (TNFα) and Shield-1 for 18 hours before sorting for mCherry. Isoclonal and polyclonal populations were created as described (32). Sorting and analysis of cells infected was performed on a FACSAria II™. Inducible-Tat cells were generated by transducing Jurkat cells with Tet-Tat-Dendra-FKBP and SFFV-rTta lentivirus at high MOI (27). The cells were incubated in Dox for 24 h and then FACS sorted for Dendra+ cells to create a polyclonal population. To create the Tet-Tat-Dendra-FKBP + LTR-mCherry cells, the polyclonal population was infected with LTR-mCherry lentivirus at a low MOI. Before sorting for mCherry+ and Dendra+ cells, Dox was added at 500ng/mL for 24 h, and single cells were FACS sorted and expanded to isolate isoclonal populations.

### Flow Cytometry Analysis

Flow cytometry data was collected on a BD FACSCalibur™ DxP8, BD LSR II™, or HTFC Intellicyt™ for stably transduced lines and sorting. Flow cytometry data was analyzed in FlowJo™ (Treestar, Ashland, Oregon) and using customized MATLAB® code (27).

### Mathematical Model and Stochastic Simulations

A simplified two-state model of LTR toggling and Tat positive feedback was constructed based on experimental data of LTR toggling (24,36) and simulated using the Gillespie algorithm (55) in MATLAB® to test how altering toggling kinetics and feedback strength would affect the activity of the circuit. At least 1000 simulations were run for each condition.

Alternatively to sweep the parameter space of different modulations of the Tat circuit, the <u>a</u>ccurate <u>c</u>hemical <u>m</u>aster <u>e</u>quation (ACME) method (56,57) was used to directly solve the chemical master equation (CME) to obtain the full probability landscapes of protein copy number. For each parameter pair in the sweeping, the protein probability landscape was computed at day 3 or at steady state. The phenotype of bimodality or unimodality at different parameter pairs was based on the numbers and locations of probability peaks in the landscape using the bimodality analysis approach described in the methods below.

### Bimodal Analysis

Two approaches were taken to quantify whether a distribution from the experimental data or simulations was bimodal or unimodal. The first, applied to both simulations and experimental data, was to convert the fluorescence density data using the *bkde* function in the KernSmooth package in R to a binned kernel density (58): the KernSmooth R package is available at https://cran.rproject.org/web/packages/KernSmooth/index.html. To filter out biologically irrelevant noise in the data, the data points with fluorescence density less than 1 or small peaks lower than 0.05 in calculated kernel density function were ignored. The number of modality peaks was performed by calculating the second order derivative of the kernel density. The second approach, only applied to the experimental data, was to utilize the Hartigan Dip Test, a dip statistic testing for multimodality by testing for maximal differences which can ascertain the probability that a particular distribution is unimodal (39). Code for the Hartigan Dip Test was obtained from http://nicprice.net/diptest/, adapted from Hartigan’s original Fortran Code for Matlab®.

## ACKNOWLEDGEMENTS

We thank members of the Weinberger, and Simpson Laboratories, as well as Hana El-Samad for helpful discussions. We especially thank Charles Chin and Adrian Jacobo for help with the stochastic simulations.

## SUPPORTING INFORMATION CAPTIONS

**<u>Table S1.</u> Chemical reaction scheme (with parameters) for stochastic simulations of circuits where Tat only modulates k_OFF_.** For the open-loop circuit with no positive feedback, the ** reaction is present, but the * reaction is not. The variable input in the ** parameter represents the different Tat inputs, which experimentally are varied by adding different amounts of doxycycline to the culture (Fig. 2). The * reaction closes the loop and is used to model the circuit that has positive feedback. The ** reaction is not used for the models with positive feedback. The fifth reaction defines Tat’s modulation of promoter toggling. For Tat affecting k_OFF_, Tat binds to the LTR_ON_ state and creates a third state, LTR_Tat_. From the LTR_Tat_ state, the LTR must move through LTR_ON_ first before switching back ‘OFF’.

**<u>Table S2.</u> Chemical reaction scheme (with parameters) for stochastic simulations of circuits where Tat only modulates k_ON_.** See Table S1 description for further information. The fifth reaction represents Tat’s ability to modulate burst frequency through k_ON_. Tat binds to the LTR_OFF_ state, and flips the promoter to the LTR_ON_ state.

**<u>Table S3.</u> Chemical reaction scheme (with parameters) for stochastic simulations of circuits where Tat only modulates alpha.** See Table S1 description for further information. The fifth reaction defines Tat’s modulation of burst size by modulating alpha. Tat binds to the LTR_ON_ state and promotes transcription, thereby affecting transcriptional efficiency when the promoter is already in an active state.

**<u>Figure S1.</u> Promoter toggling kinetics controls the separation of gene-expression peaks due to transient production and decay.** For a given time, the rate of switching between the ‘ON’ and ‘OFF’ promoter state (top pulse trains) is related to the duration of time in a specific promoter state. The duration of the promoter state determines the length, or separation from the mean (cyan line, same value for each panel) of the transient production or decay of gene-expression products. Increasing promoter kinetics reduces transients and the separation between potential peaks in a bursty system (top left moving to the right, then bottom left moving to the right).

**<u>Figure S2.</u> The LTR produces bimodal distributions in response to unimodal Tat inputs.** (A) Histograms of the Tat input to the LTR, as measured by Dendra fluorescent signal, is unimodal across all combinations of doxycycline and Shield-1. The colors of the lines indicate increasing doxycycline concentrations (red, 0ng/mL → orange, 2.5ng/mL → yellow, 5ng/mL → green, 12.5ng/mL → cyan, 25ng/mL → blue, 50ng/mL → pink, 250ng/mL → magenta, 500ng/mL), and the increasing brightness of the same color represents increasing Shield-1 concentrations (0, 10, 50, 100, 500, and 1000nM). (B) Histograms of LTR output as measured by mCherry fluorescent signal. The ‘Dim’/‘Bright’ threshold was set based on each population’s mCherry expression in the absence of doxycycline or Shield-1 (i.e. no Tat). The change in signal in the ‘Bright’ population is used to determine the small-signal loop gain (Fig. S11) in response to Tat.

**<u>Figure S3.</u> Bimodality analysis for the open-loop system.** 9 isolconal populations of the open-loop circuits described in Figure 2 were exposed to 48 different doxycycline or Shield-1 concentrations. The populations were assessed for the number of modes as described in the methods. Briefly, fluorescence intensity data was smoothed using the *bkde* function in the KernSmooth package in R to a binned kernel density. The number of modality peaks was calculated by taking the second order derivative of the kernel density. Gray squares are unimodal and black squares are bimodal.

**<u>Figure S4.</u> Tat activation of the LTR controls expression pulses.** Single-cell time-lapse fluorescence microscopy of the open-loop circuit without doxycycline (black lines) or with 25ng/mL (red lines), 100ng/mL (cyan lines) or 500ng/mL (green lines) of doxycycline. Both Dendra (i.e. Tat levels) and mCherry (i.e. LTR activity) fluorescence levels were tracked over time. Variable Tat inputs as measured by Dendra fluorescence leads to variable expression pulses from the LTR as measured by Cherry expression.

**<u>Figure S5.</u> Full-length HIV decision-making can be controlled in the absence of feedback or cooperativity.** Schematic of the full-length HIV open-loop circuit (top). Doxycycline addition induces Tat expression, which can activate expression of the full-length HIV virus with a fluorescent mCherry reporter. Cells were initially infected in the absence (red histogram) or presence of doxycycline (blue histogram), and a time point was taken 24hrs post infection (left side, ‘Initial Infection’). Doxycycline was then added to a split of the ‘No Dox’ at the Initial Infection for 24hrs to look for HIV reaction (right side, ‘Latent Reactivation’).

**<u>Figure S6.</u> The fluctuations in mCherry depend on the mechanism of Tat transactivation.** (A) Each parameter set was allowed to run for 1000 stochastic simulations, where Tat would work through k_ON_ (green), k_OFF_ (pink), or alpha (black lined) alone. The average protein count is equivalent for all the simulations. (B) Time course of mCherry count over time, showing the extent of stochastic fluctuations when Tat affects k_ON_, k_OFF_, or alpha. Three representative traces are shown for each.

**<u>Figure S7.</u> Bimodality analysis for the closed-loop feedback system.** 9 isoclonal populations were exposed to various concentrations of Shield-1 as described in Figure 3. The number of modes was determined as described in the methods. Briefly, fluorescence intensity data was smoothed using the *bkde* function in the KernSmooth package in R to a binned kernel density. The number of modality peaks was calculated by taking the second order derivative of the kernel density. Gray squares are unimodal and black squares are bimodal.

**<u>Figure S8.</u> Positive feedback strength sets the steady-state activity and percentage of cells in an active state.** A polyclonal population of Ld2GITF cells were exposed to TNF-alpha for 24 hours (-24 to 0hrs), then the cells were washed and split into two cultures, one with Shield-1 (1uM, blue) or one in the absence of Shield-1 (0uM, gray). GFP measurements were taken every 24 hours and mean fluorescence intensity (right axis) or the percentage of cells in the ON state (left axis) were quantified. In the absence of Shield-1 after 72hrs, the cells returned to the unperturbed state in both percent ON and mean fluorescence intensity. In the presence of Shield-1, positive feedback strength is increased, and the system remains activated for a longer duration of time. Importantly, both populations return to the state of no TNF-alpha addition, i.e. no bistability.

**<u>Figure S9.</u> Simulations of Tat modulating two parameters of transcriptional bursting.** We consider three different phenotypes: unimodality of latency (blue areas), unimodality of activation (red areas), and bimodality (yellow areas). The phase diagrams of phenotypes for three different modulations: k_ON_-k_OFF_ (left graphs), k_ON_-alpha (middle graphs), and k_OFF_-alpha (right graphs) based on the steady state probability landscapes are shown in part A, and the phenotype phase diagrams based on the day 3 probability landscapes are shown in part B. Details about the models and parameter sweeping can be found in the Materials and Methods. In the modulations of k_ON_-k_OFF_ (left graphs) and k_ON_-alpha (middle graphs), some parameter pairs are bimodal at day 3 (yellow area in part B), but become unimodality of activation at the steady state (red area in part A). This is due to the slow evolution of the probability landscape in these parameter pairs. The phenotypes of all parameter pairs in k_OFF_-alpha (right graphs) modulation at steady state are consistent with those at day 3. All simulations were started with initial toggling kinetics of k_ON_ = 0.001/min, k_OFF_ = 0.01/min, and the rest of the parameters can be found in Table S1.

**<u>Figure S10.</u> Tat positive feedback is non-latching.** (A) A schematic showing the input-output relationship for a positive feedback loop under the control of a constitutive promoter. Unimodal signal inputs of varying strengths reach a constitutive promoter encoding for a transcription factor (TF), which initiates positive feedback. The level of amplification due to positive feedback is quantified by the small-signal loop gain. For loop gains < 1 across all protein concentrations, the system displays non-latching feedback and the results in a unimodal output over the abundance regime. However, if small-signal loop gain increases with protein abundance to ~1, small input fluctuations are drastically amplified and can generate a bimodal distribution in the output (bottom right). The error bars around the circles in A, right-hand graphs, represent, for a population of cells that receive the same inputs, the fluctuations that would lead some cells to display higher or lower small-signal loop gains. (B) Quantification of the small-signal loop gain for the closed-loop circuit in nine isoclonal populations reveals that Tat feedback is non-latching.

**<u>Figure S11.</u> Quantification of the open-loop small-signal loop gain shows non-latching feedback.** (A) Plot of the fold change in Tat-Dendra abundance versus the fold-change in mCherry ‘ON’ population expression for nine isoclonal populations. (B) Quantification of the small-signal open-loop gain of the nine isoclonal populations. These values are representative of the expected small-signal loop gain for an intact circuit with feedback. Importantly, all nine isoclonal populations indicate that Tat feedback is non-latching.

**<u>Figure S12.</u> The LTR’s response to Tat is biphasic; the LTR is sensitive to low levels of Tat, but insensitive to higher levels of Tat.** Plot of normalized LTR-mCherry output to normalized Tat-Dendra fluorescence for the eleven clonal populations (Figure 2). The data is best fit with a logarithmic function but can also be represented with two linear fits (R^2^ = 0.98): one for the sensitive region (between 0 and 0.2, Normalized Tat-Dendra Fluorescence) and insensitive region (between 0.2 and 1, Normalized Tat-Dendra Fluorescence).

**<u>Figure S13.</u> Non-monotonic relation between the percent of toggling kinetics that yield bimodality versus feedback strength.** To simulate various feedback strengths the binding affinity, k_b_, was tuned from 5x10^-7 to 10, on a parameter scan across k_ON_ and k_OFF_ values ranging from 0.001 to 10/minute. Next, of those parameter scans, a bimodality test was performed (see the Bimodality Analysis in the Methods). The percentage of parameters that yielded bimodality was then quantified. Various thresholds were set to determine whether a population was bimodal by requiring that each mode had to have 10^-8%, 0.1%, 1%, 5%, or 10% of the total population.

